# Trans-Endothelial Insulin Transport is Impaired in Skeletal Muscle Capillaries of Obese Male Mice

**DOI:** 10.1101/585372

**Authors:** Ian M Williams, P Mason McClatchey, Deanna P Bracy, Jeffrey S Bonner, Francisco A Valenzuela, David H Wasserman

**Affiliations:** Department of Molecular Physiology and Biophysics, Vanderbilt University, Nashville, Tennessee, USA; Mouse Metabolic Phenotyping Center, Vanderbilt University, Nashville, Tennessee, USA; Lilly Research Laboratories, Indianapolis, Indiana, USA

## Abstract

Delivery of insulin to the surface of myocytes is required for skeletal muscle (SkM) insulin action. Previous studies have shown that SkM insulin delivery is reduced in the setting of obesity and insulin resistance (IR). The key variables that control SkM insulin delivery are 1) microvascular perfusion and 2) the rate at which insulin moves across the continuous endothelium of SkM capillaries. Obesity and IR are associated with reduced insulin-stimulated SkM perfusion. Whether an impairment in trans-endothelial insulin transport (EIT) contributes to SkM IR, however, is unknown. We hypothesized that EIT would be delayed in a mouse model of diet-induced obesity (DIO) and IR. Using intravital insulin imaging, we found that DIO male mice have a ~15% reduction in EIT compared to their lean counterparts. This impairment in EIT is associated with a 45% reduction in the density of endothelial vesicles. Despite impaired EIT, hyperinsulinemia sustained delivery of insulin to the interstitial space in DIO male mice. Even with maintained interstitial insulin delivery DIO male mice still showed SkM IR, indicating severe myocyellular IR in this model. Interestingly, there was no difference in EIT, endothelial ultrastructure or SkM insulin sensitivity between lean and high fat diet-fed female mice. These results suggest that, in male mice, obesity results in damage to the capillary endothelium which limits the capacity for EIT.

## INTRODUCTION

Skeletal muscle (SkM) comprises the vast majority of insulin-sensitive tissue. As such, it plays a key role in insulin-stimulated glucose disposal. Furthermore, SkM is a major site of insulin resistance (IR) during obesity (1, 2) and SkM IR precedes the development of Type 2 diabetes (3). Determining the mechanism of SkM IR, therefore, is critical to developing therapies that restore metabolic function in obese individuals.

Insulin action in SkM depends on the delivery of insulin to the myocyte, binding of insulin to the insulin receptor, and subsequent activation of signaling cascades. Following its appearance in the systemic circulation, insulin stimulates endothelium-dependent relaxation of SkM arteries and arterioles (4). This vasodilation enhances SkM perfusion and is thought to increase the capillary surface area available for glucose and insulin exchange in SkM (5). Subsequently, insulin must transit the continuous endothelium of SkM capillaries by a nonsaturable, fluid-phase transport process (6, 7). The endothelium is a significant barrier which restricts insulin access to SkM and, thus, is a key regulator of insulin action in muscle (8, 9). Once insulin accesses the interstitial space, it binds the insulin receptor on myocytes and stimulates the insulin receptor - phosphoinositide 3-kinase (PI3K) / Akt signaling cascade (10). Ultimately, these signaling events induce translocation of glucose transporter type 4 (Glut4) to the plasma membrane of SkM, thereby increasing glucose transport (11).

Many investigators have observed a reduction in SkM insulin signaling (reviewed in Ref 12) and glucose transport (13, 14) in the setting of obesity, IR, and type 2 diabetes. This has generally been attributed to the toxic lipid accumulation (15), mitochondrial dysfunction (16), or inflammation (17) that occurs in obese individuals. It has also been hypothesized that decreased insulin signaling is caused by a reduction in microvascular delivery of insulin to myocytes (9). That is, pathological processes occurring outside the myocyte may contribute to SkM IR.

Most (18–20), but not all (21), previous studies have found that the delivery of insulin to the interstitial space in SkM is reduced in obese humans and animal models. This observation of impaired interstitial insulin delivery suggests that microvascular dysfunction may contribute to SkM IR. The delivery of insulin to SkM is determined by 1) the surface area available for insulin exchange and 2) the rate at which insulin moves across the capillary endothelium. Several studies have demonstrated that insulin-stimulated SkM perfusion is reduced in obese, insulin resistant, and type 2 diabetic humans (as reviewed in Ref 5). This decrease in perfusion would be expected to reduce the surface area for insulin exchange and, therefore, microvascular insulin delivery. Whether the rate of trans-endothelial insulin transport (EIT) is altered in obesity and IR, however, is unknown.

We hypothesized that EIT would be reduced in the setting of obesity and SkM IR. To address this hypothesis, we utilized the high-fat diet (HFD)-fed mouse model of obesity and IR (22). We found that HFD-fed male mice had a 45% reduction in the density of capillary endothelial vesicles, the putative vehicles for EIT. This abnormal capillary endothelial ultrastructure was associated with a 15% reduction in EIT, as determined by quantitative intravital microscopy (6). Although EIT was reduced, we still observed significant impairment of insulin sensitivity at the level of the myocyte. Conversely, female mice were protected from the effects of HFD on adiposity, endothelial ultrastructure, EIT, and SkM insulin sensitivity. These findings indicate that HFD results in abnormal capillary endothelial ultrastructure and impaired EIT in male mice.

## RESEARCH DESIGN AND METHODS

### Mouse models

Male and female 6-week old C57Bl/6J mice were randomized to either a standard chow (5001 Laboratory Rodent Diet; Lab Diet) or HFD (60% calories from fat; Bioserv F3282) for 16 weeks. We chose this length of HFD because it induces SkM IR (23). Mice were purchased from Jackson Laboratory (Bar Harbor, ME, USA) and bred and housed either at Vanderbilt University or Jackson Laboratory. One HFD-fed male mouse which underwent intravital microscopy was excluded due to a lack of weight gain and fat accumulation. Mice were housed in cages containing Pure-o’Cel^®^ (The Andersons Inc.; Maumee, OH, USA) paper bedding with 0 to 4 littermates and free access to food and water. Cages were kept in a pathogen-free, temperature and humidity-controlled facility and maintained on a 12-hour light dark cycle. All animal procedures were approved by the Vanderbilt Institutional Animal Care and Use Committee and conducted in accordance with the NIH Guide for the Care and Use of Laboratory Animals.

### Body composition

Body composition was determined after 15 weeks of chow or HFD using a Minispec mq10 NMR analyzer (Bruker; Billerica, MA, USA).

### Electron microscopy

Samples were processed for transmission electron microscopy and imaged in the Vanderbilt Cell Imaging Shared Resource – Research Electron Microscopy facility. Mice were anesthetized with sodium pentobarbital and tissues were fixed by perfusing mice with 2.5% glutaraldehyde in 0.1mol/l cacodylate buffer through a catheter inserted into the left ventricle. The gastrocnemius was then excised and fixed for an additional 24h at 4°C. Tissues were subsequently postfixed with osmium tetroxide, dehydrated and embedded in Epon 812 epoxy resin. Ultrathin sections (70-80nm) were cut from embedded tissues and stained with 2% uranyl acetate and Reynold’s lead citrate. Sections were imaged using a Philips/FEI Tecnai T12 electron microscope (ThermoFisher Scientific).

Capillaries were identified and selected for analysis at low magnification where endothelial ultrastructure (i.e. vesicles) is not visible. This approach prevents bias in capillary selection. Subsequently, 2-4 images were acquired of each capillary at high magnification (67,000x). 6-13 capillaries were imaged per section. Endothelial vesicular density was measured by applying a test point grid (ImageJ; NIH) to micrographs and then counting the number points residing within vesicles and within the entire endothelium (24). Endothelial vesicular density was calculated as the number of points within vesicles relative to the number of points in the endothelium as a whole. Vesicular diameter was measured at the widest point of each vesicle. Endothelial vesicle localization was assessed by manually counting vesicles that were either facing the capillary lumen (luminal), contained within the endothelium (intracellular), or facing the interstitial space (abluminal). Basement membrane thickness was measured perpendicularly to points of intersection between a line grid applied to micrographs and the capillary endothelium.

### Catheterization

Catheterization of the carotid artery and jugular vein was performed as described previously (25, 26). For intravital microscopy experiments, only jugular vein catheters were implanted. For insulin tolerance tests, both the carotid artery and jugular vein were catheterized. Mice were allowed to recover for at least 3 days following jugular vein catheterizations and at least 5 days following combined carotid artery and jugular vein catheterization. Recovery from catheterization surgeries was deemed adequate if mice lost no more than 10% of their preoperative body weight and if their behavior was normal.

### Intravital microscopy

*In vivo* imaging of a fully bioactive fluorescent insulin (INS-647) probe was performed as described previously in great detail (6). Briefly, mice were placed in plastic buckets containing Pure-o’Cel^®^ paper bedding and food and water were removed for 5 hours beginning in the morning and ending in the early afternoon. Mice from different sex and diet groups were studied in a random, alternating temporal order on a given day and then this order was systematically shifted on subsequent days to prevent time-of-day effects. After the 5-hour fast, mice were anesthetized by administering a ketamine/xylazine/acepromazine cocktail (KXA; 7.9/1.6/0.2 mg/kg) through the indwelling jugular vein catheter. We chose to use KXA because it minimizes mouse motion, maintains gastrocnemius blood flow, and is easy to administer. The lateral gastrocnemius was then exposed by carefully removing the overlaying skin and fascia. Next, mice were positioned on a custom stage mount and the exposed gastrocnemius was continuously superfused with a bicarbonate-buffered (18mM) physiological saline solution (132mmol/l NaCl, 4.7mmol/l KCl, 2mmol/l MgSO4, 1.2mmol/l CaCk) which was maintained at 37°C and pH 7.4. Body temperature was monitored using a rectal probe and maintained with an electric blanket connected to a homeothermic feedback controller system (Harvard Apparatus; Cambridge, MA, USA). The lateral gastrocnemius was imaged using a 20x 0.8NA Plan-Apochromat air objective on an LSM 780 confocal microscope (Zeiss; Oberkochen, Germany). After ensuring the quality of the tissue preparation, mice received a 50μg bolus (in 50μL 0.9% NaCl) of a 2 megadalton tetramethylrhodamine-labeled dextran (rho-dex; Thermo Fisher Scientific; Waltham, MA, USA). Rho-dex is too large to leak from SkM capillaries and therefore serves as a vascular marker.

After selecting a field of view and acquiring a background image, mice received a 4U/kg body mass bolus of INS-647 and 13μCi (0.481MBq) of 2[^14^C]deoxyglucose (2[^14^C]DG; specific activity: 9.25-13.0GBq/mmol; Perkin Elmer; Waltham, MA, USA) through the indwelling jugular vein catheter. 4-slice z-stacks (spaced 4μm apart) of the rho-dex and INS-647 channels were acquired every minute for 15 minutes following the INS-647 bolus. The confocal pinhole was set to give optical sections of 8μm (+/- 4μm about the focal plane). Following the imaging experiment, blood glucose was measured from the tail vein (Accu-Chek; Roche; Basel, Switzerland), mice were sacrificed by cervical dislocation, and tissues were harvested for analysis. For albumin imaging experiments, mice received a 5mg/kg bolus of Alexa Fluor 647-labeled bovine serum albumin (Alb-647; ThermoFisher Scientific). Subsequently, Alb-647 imaging was performed identically to INS-647 imaging.

For venular imaging, we initially located venules based on their size, high permeability, and timing of blood flow arrival. 2 or 4-slice z-stacks (spaced 5μm apart) of rho-dex and INS-647 in venules were acquired with a 10X 0.5NA Fluar air objective. The confocal pinhole was set to give optical sections of 10μm (+/- 5μm about the focal plane). As with capillary imaging, images of rho-dex and INS-647 channels were acquired prior to the INS-647 bolus and every minute for 15 minutes thereafter.

### Image analysis

Prior to quantification, images were inspected visually to check for major motion artifacts, significant rho-dex leakage, or fields of view with very few capillary segments. Mice or individual frames which did not meet these criteria were excluded from subsequent analysis. Images were segmented using an automated Otsu-based thresholding algorithm to demarcate capillaries and the interstitial space (6). The interstitial space was defined as the region emanating radially 1-3μm from the capillary wall. INS-647 concentrations were measured in the capillary and interstitial space as a function of time following the INS-647 bolus. We measured extravascular INS-647 by dilating the vascular mask by 1μm and inverting it to segment the entire extravascular space in the field of view.

### Image quantification

The clearance of INS-647 from the plasma was analyzed by fitting the first 5 minutes of the plasma INS-647 decay curve to a mono-exponential decay function. We decided to describe plasma INS-647 with mono-exponential decay as our imaging data fit this function better than bi-exponential decay. The rate of EIT can be assessed by determining the rate at which insulin in the capillary equilibrates with the surrounding interstitial space. To measure this equilibration rate, we plotted the ratio of plasma to interstitial INS-647 intensity as a function of time following the INS-647 bolus. Over time, this ratio will decay as INS-647 leaves the plasma and enters the interstitial space. We quantified the gradient decay constant of the plasma/interstitial INS-647 ratio by fitting the first 5 minutes of the decay curve to a mono-exponential function. Plasma-perfused surface area was determined using the rho-dex (vascular marker) channel. Capillary diameter was estimated from rho-dex images by calculating the ratio of vascular surface area to vascular perimeter. This ratio approximates the radius of long cylindrical vessel segments (27).

### Insulin tolerance tests

To test whether SkM IR could be detected in DIO male mice, we performed intravenous insulin tolerance tests using the same protocol as intravital microscopy (i.e. anesthetized, 4U/kg insulin tolerance tests). The objective of these experiments was to have measurements of insulin sensitivity that would parallel the results obtained during intravital microscopy. Following a 5-hour fast (see **Intravital Microscopy**) mice were anesthetized with KXA and allowed to stabilize for 10 minutes. Following a baseline arterial glucose sample, mice received a combined bolus of 4U/kg insulin (Novolin R; Novo Nordisk; Bagsværd, Denmark) and 13μCi 2[^14^C]DG. Arterial glucose was measured from whole blood at t = 2, 5, 10, 15, 20, and 30 min. Additional plasma samples were collected at these time points for measurement of plasma 2[^14^C]DG. At t = 30 min, mice were sacrificed by cervical dislocation and tissues were harvested for analysis.

### Measurement and analysis of isotopic glucose accumulation

Plasma 2[^14^C]DG and tissue 2[^14^C]DG-6-phosphate (2[^14^C]DGP) were measured as described previously (28). Tissue glucose clearance (Kg; μL•g tissue^-1^ • min ^-1^) was calculated using the following equation:

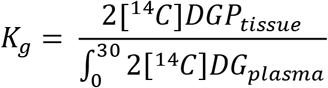

where 2[^14^C]DGP_tissue_ is the accumulated 2[^14^C]DGP in the tissue (dpm•g^-1^ tissue) and 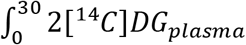 is the time-integrated 2[^14^C]DG activity in the plasma calculated using trapezoidal integration.

### Single-cell RNA sequencing

Measurements of insulin receptor mRNA levels in single endothelial cells (ECs) from various mouse tissues were obtained using the publicly available *Tabula Muris* dataset (29). These measurements were obtained from 4 male and 3 male 10-15 week old C57Bl/6JN mice.

### Statistics

Statistical analysis was performed using Prism 7 (Graphpad; San Diego, CA, USA). Grubbs’ test was performed to detect individual outliers. No statistical outliers were detected or removed. Data are presented as mean ± standard error of the mean. Groups were compared using either unpaired Student’s t-tests or two-way repeated measures ANOVA.

## RESULTS

### HFD causes ultrastructural defects in the SkM capillary endothelium of male mice

Obesity and SkM IR are strongly associated with endothelial dysfunction (30). To determine whether this dysfunction may affect EIT, we initially characterized capillary endothelial ultrastructure in a mouse model of diet-induced obesity (DIO) and SkM IR. Lean and DIO mouse models were generated by feeding male mice chow or HFD for 16 weeks. As expected, DIO mice weighed 52% more than their lean counterparts (**Supplemental Figure 1A**) and had a 2.5-fold increase in body fat percentage (**Supplemental Figure 1B**).

Using transmission electron microscopy, we observed that the density of vesicles in the endothelium was reduced by ~45% in DIO male mice (**Figure 1A-C**). Neither the diameter of vesicles (**Figure 1D**), nor the localization of vesicles within the endothelium (**Figure 1E**), however, were altered in DIO male mice. Furthermore, the basement membrane thickness was not different between groups (**Figure 1F**). This observation of reduced endothelial vesicles, which can serve as either distinct vesicular transporters or form trans-endothelial channels (31), raises the possibility that EIT is impaired in DIO male mice.

**Figure 1:**
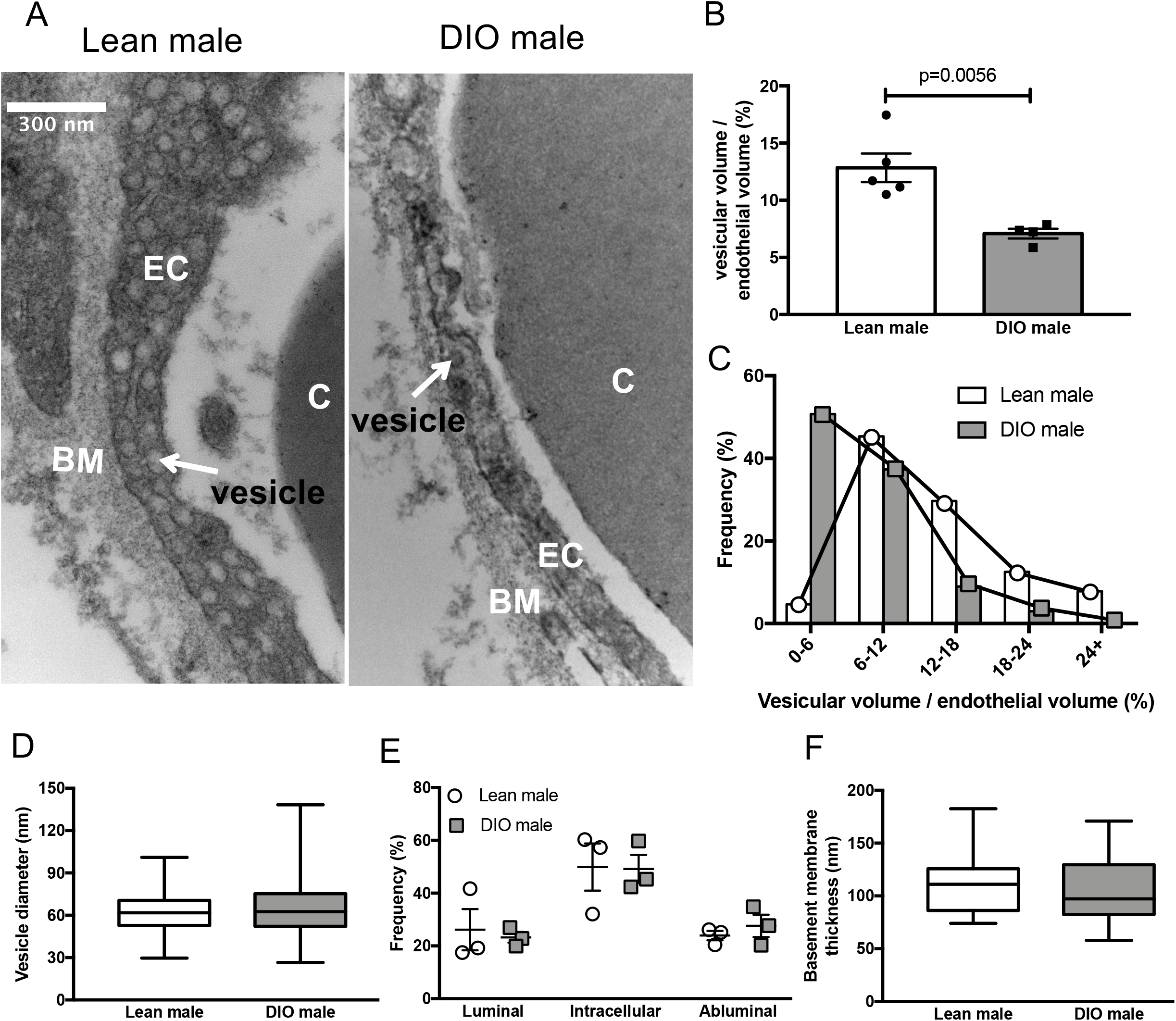
Skeletal muscle capillaries of DIO male mice contain fewer endothelial vesicles. **A)** Representative electron micrographs of the capillary endothelium in the gastrocnemius of lean and DIO male mice. **B)** Volume of vesicles relative to total endothelial volume in lean (n=5) and DIO (n=4) male mice. **C)** Frequency distribution of relative vesicular volume in all capillaries grouped from lean (n=64) and DIO male mice (n=66). **D)** The average diameter of all endothelial vesicles in lean (n=227) and DIO (n=116) male mice. **E)** Frequency distribution of the localization of vesicles in the capillary endothelium. **F)** Average basement membrane thickness in capillaries from lean (n=31) and DIO male mice (n=31). In the box and whisker blots, the box extends from the 25^th^ to the 75^th^ percentiles and the whiskers indicate the range. Groups were compared using Student’s t-test. C – capillary lumen, EC – endothelial cell, BM – basement membrane.

### HFD impairs trans-endothelial insulin transport in male mice

Given that DIO male mice showed a reduction in endothelial vesicles, putative vehicles for EIT, we hypothesized that EIT would be similarly reduced in DIO male mice. Using intravital insulin imaging, we observed that EIT is impaired in DIO male mice (**Figure 2**). Namely, the rate at which plasma INS-647 equilibrated with the interstitial space was ~15% lower in DIO mice compared to lean mice (**Figure 2B&C**). This indicates that EIT is slower in DIO male mice. Following the INS-647 bolus, capillary INS-647 levels were consistently higher in DIO male mice (**Figure 3A**). The clearance rate of plasma INS-647, however, was not significantly different between groups (**Figure 3A&B**). Consistent with higher levels of plasma INS-647, DIO male mice also had higher absolute levels of interstitial INS-647 (**Figure 3C**). The rate of interstitial INS-647 appearance, however, was slightly, but non-significantly, lower in the DIO male mice (**Figure 3D**). Furthermore, the magnitude of the increased interstitial INS-647 in DIO male mice (**Figure 3C**) was lower than that of the increase in plasma INS-647 (**Figure 3A**). These findings indicate that, while DIO male mice had higher levels of plasma INS-647, a lower fraction of the plasma INS-647 was able to access the interstitial space. This demonstrates impaired EIT.

**Figure 2:**
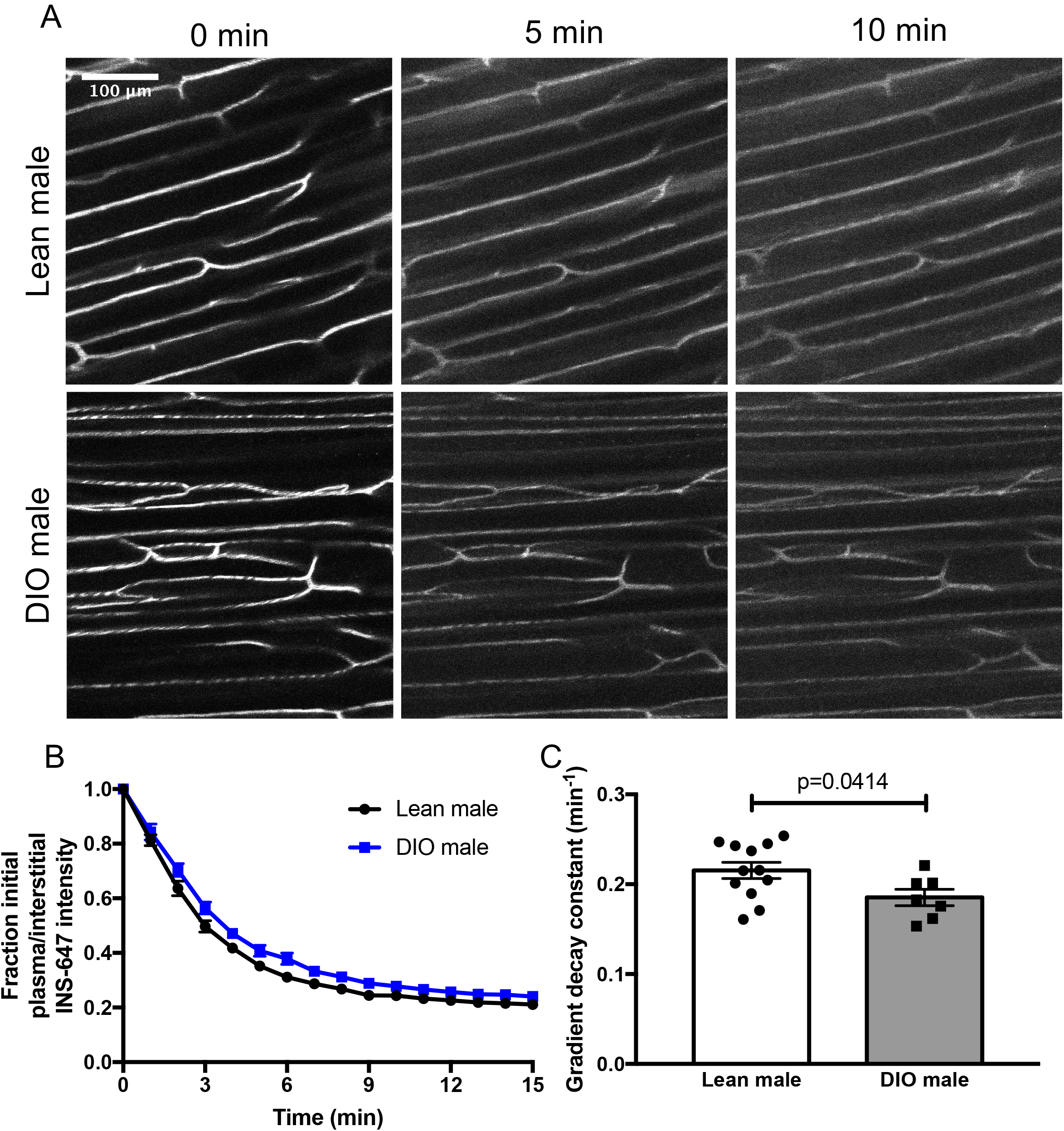
Obese male mice have impaired trans-endothelial insulin transport in skeletal muscle capillaries. **A)** Representative INS-647 images (maximum intensity projections) in lean (n=12) and DIO (n=7) male mice. **B)** The ratio of plasma to interstitial INS-647 as a function of time following INS-647 injection, normalized to the ratio at t = 0 min. **C)** Decay constant of the plasma / interstitial INS-647 ratio, a measure of trans-endothelial insulin transport kinetics. INS-647 – insulin-647, DIO – diet-induced obese. Groups were compared using Student’s t-test.

**Figure 3:**
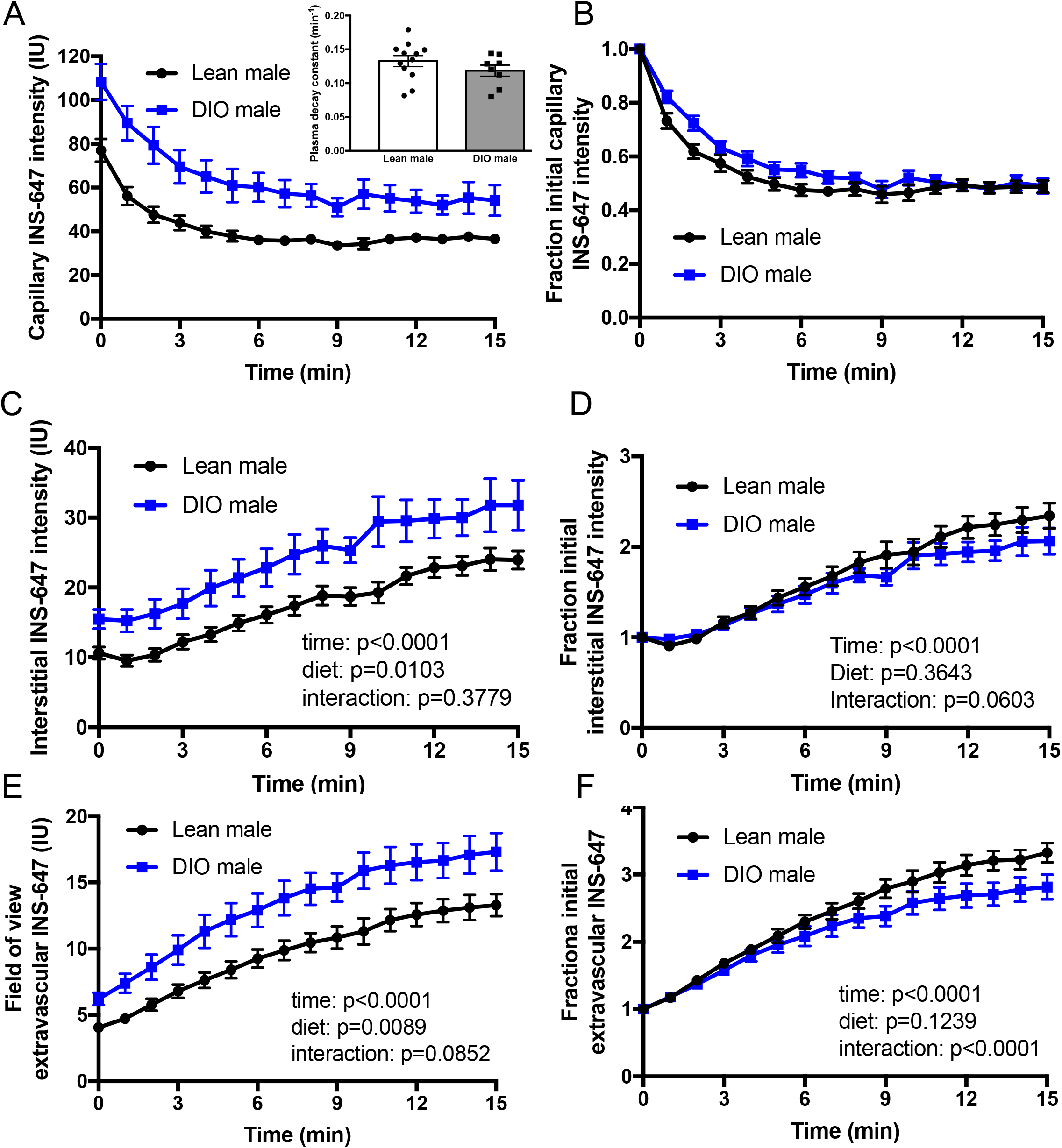
Effects of HFD on plasma insulin clearance, interstitial appearance, and insulin delivery in male mice. **A)** Capillary plasma INS-647 intensity as a function of time following injection in lean (n=12) and DIO (n=7) male mice. Inset shows the decay constant of capillary INS-647. **B)** Data in **A** normalized to the capillary INS-647 intensity at t = 0 min. **C)** Interstitial INS-647 intensity as a function of time following injection. The interstitial space is defined as the region emanating 1-3μm from the capillary wall. **D)** Data in **C** normalized to the interstitial INS-647 intensity at t = 0 min. **E)** Total extravascular INS-647 in the field of view as a function of time following injection. **F)** Data in **E** normalized to the extravascular INS-647 intensity at t = 0 min. Groups were compared either by Student’s t-test or by two-way repeated measures ANOVA. IU – intensity units.

EIT is one of three variables, including plasma insulin concentration and the capillary surface area for insulin exchange, that control total SkM insulin delivery. We found that neither plasma-perfused surface area (**Supplemental Figure 2A&B**) nor capillary diameter (**Supplemental Figure 2C&D**) were affected by HFD. This suggests that the surface area for insulin exchange is the same in lean and DIO males. To determine whether the reduced EIT in DIO males affected total insulin delivery, we measured INS-647 in the entire extravascular space. We found that, while the extravascular INS-647 levels were consistently higher in DIO male mice (**Figure 2E**), the rate of INS-647 appearance in the extravascular space was reduced (**Figure 2F**). These findings indicate that the kinetics, but not absolute levels, of INS-647 delivery to SkM are impaired in DIO male mice.

Following intravital microscopy experiments, we observed that DIO male mice had higher blood glucose levels (**Supplemental Figure 3A**) and lower accumulation of the isotopic glucose tracer 2[^14^C]deoxyglucose-6-phosphate (2[^14^C]DGP) in the gastrocnemius (**Supplemental Figure 3B**). These findings indicate that DIO male mice have whole-body and SkM IR. To confidently ensure that SkM IR could be detected using the same conditions as intravital microscopy experiments, we performed insulin tolerance tests with isotopic glucose tracers in a separate cohort of DIO male mice (**Figure 4**). As expected, DIO male mice had a significantly impaired glucose-lowering response to insulin (**Figure 4A&B**). Furthermore, the clearance of 2[^14^C]deoxyglucose (2[^14^C]DG) by the soleus, gastrocnemius, and vastus muscles was reduced in DIO male mice (**Figure 4C**). These results confirm that DIO male mice display SkM IR. Therefore, SkM IR is associated with impaired EIT in DIO male mice.

**Figure 4:**
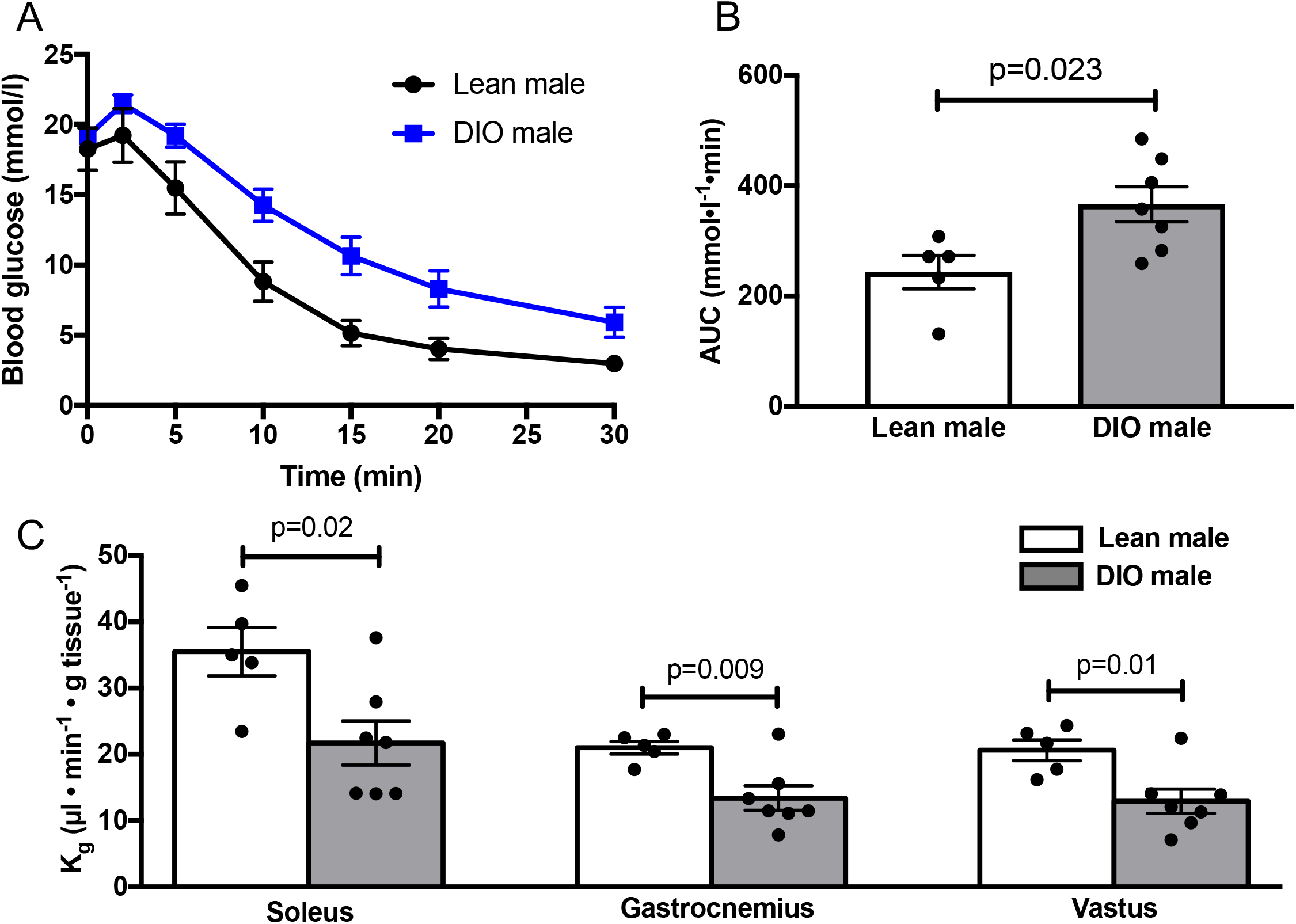
DIO male mice display skeletal muscle insulin resistance. **A)** Glucose excursions in anesthetized lean (n=5) and DIO (n=7) male mice following a 4U/kg intravenous insulin bolus. **B)** Area under the glucose excursion curves in A. **C)** Clearance of 2[^14^C]2deoxyglucose during the insulin tolerance tests by the soleus, gastrocnemius, and vastus muscles. Groups were compared using Student’s t-test.

### Venules, but not capillaries, accumulate insulin and dextran

Our previous work demonstrated that insulin transits the capillary endothelium by a fluid-phase transport mechanism that does not involve the insulin receptor (6). To further validate this finding, we assessed the expression of insulin receptor mRNA in endothelial cells (ECs) from various mouse organs using the *Tabula Muris* single-cell RNA sequencing dataset (29). We found that the majority (84%) of SkM ECs do not express the insulin receptor (**Figure 5A**). Conversely, the insulin receptor is more highly expressed in brain and liver ECs (**Figure 5A**). The relative lack of insulin receptor expression in SkM ECs confirms that insulin does not cross SkM capillary endothelium by insulin receptor-mediated transport.

**Figure 5:**
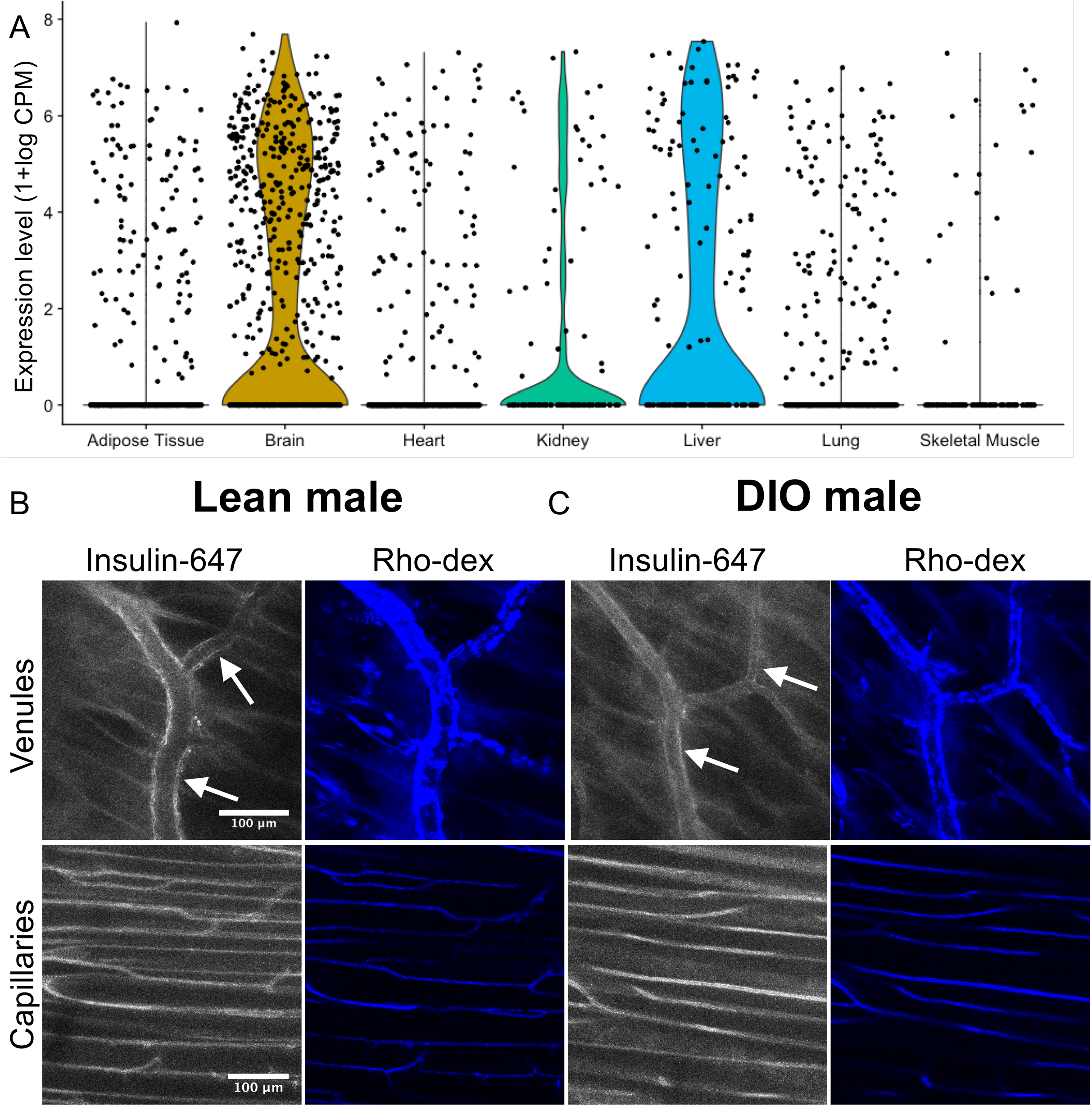
Capillary trans-endothelial insulin transport in skeletal muscle does not involve the insulin receptor or endothelial insulin accumulation. **A)** Violin plots of insulin receptor mRNA expression in tissue-specific endothelial cells as determined by single-cell RNA sequencing. **B,C)** Intravital microscopy images of insulin-647 and rhodamine-labeled 2MDa dextran (Rho-dex) in venules (top panels) and capillaries (bottom panels) from **B)** lean and **C)** DIO male mice. Both insulin and dextran can be seen accumulating in the endothelium of venules but not capillaries. Arrows indicate regions of insulin accumulation in the venular endothelium. Rho-dex – 2MDa tetramethylrhodamine-dextran, CPM – counts per million.

Insulin receptor-independent transport may involve either non-specific vesicular pinocytosis or convective movement through inter-endothelial channels. If insulin were being trafficked across the endothelium by discrete vesicular transporters, we would expect to observe INS-647 accumulation in the lining of the endothelium. We did not detect INS-647 or rho-dex accumulation in the gastrocnemius capillary endothelium in the present study (**Figure 2A, 5B&C, and 8A**) nor in our previous work (6, 27). However, we did observe a striking accumulation of INS-647 and rho-dex in cells lining post-capillary venules in both lean and DIO male mice (**Figure 5B&C**). Thus, we can detect vascular INS-647 accumulation by intravital microscopy should it exist. Taken together, these findings suggest that, in capillaries, insulin is not transported by distinct vesicular transporters but rather moves through trans-endothelial channels. Given that trans-endothelial channels mediate the transport of insulin and that EIT is impaired in DIO male mice, we posit that the decrease in endothelial vesicles (**Figure 1B**) indicates a reduction in trans-endothelial channels available for EIT.

### No difference in trans-endothelial albumin equilibration between lean and DIO male mice

To determine whether the abnormal endothelial ultrastructure observed in DIO male mice restricts the movement of other plasma-borne molecules, we measured the trans-endothelial equilibration of albumin-647 (Alb-647; **Figure 6A**). Albumin (hydrodynamic radius ~3.5nm) is a significantly larger protein than insulin (hydrodynamic radius ~1.3nm) and is usually retained within capillaries unless the barrier function of the endothelium becomes compromised (32). In contrast to INS-647, the levels of Alb-647 following injection were stable in the plasma (**Supplemental Figure 4A&B**) and interstitial space (**Supplemental Figure 4C&D**) over the course of the 15 minute experiment. Furthermore, the levels of plasma (**Supplemental Figure 4A**) and interstitial (**Supplemental Figure 4B**) Alb-647 were not significantly different between lean and DIO male mice We did not observe any differences in the ratio of plasma to interstitial Alb-647 ratio between lean and DIO male mice (**Figure 6B&C**), indicating that capillary permeability to Alb-647 is the same between groups. These findings suggest that the abnormal endothelial ultrastructure observed in DIO male mice does not alter capillary permeability to a protein of albumin’s size.

**Figure 6:**
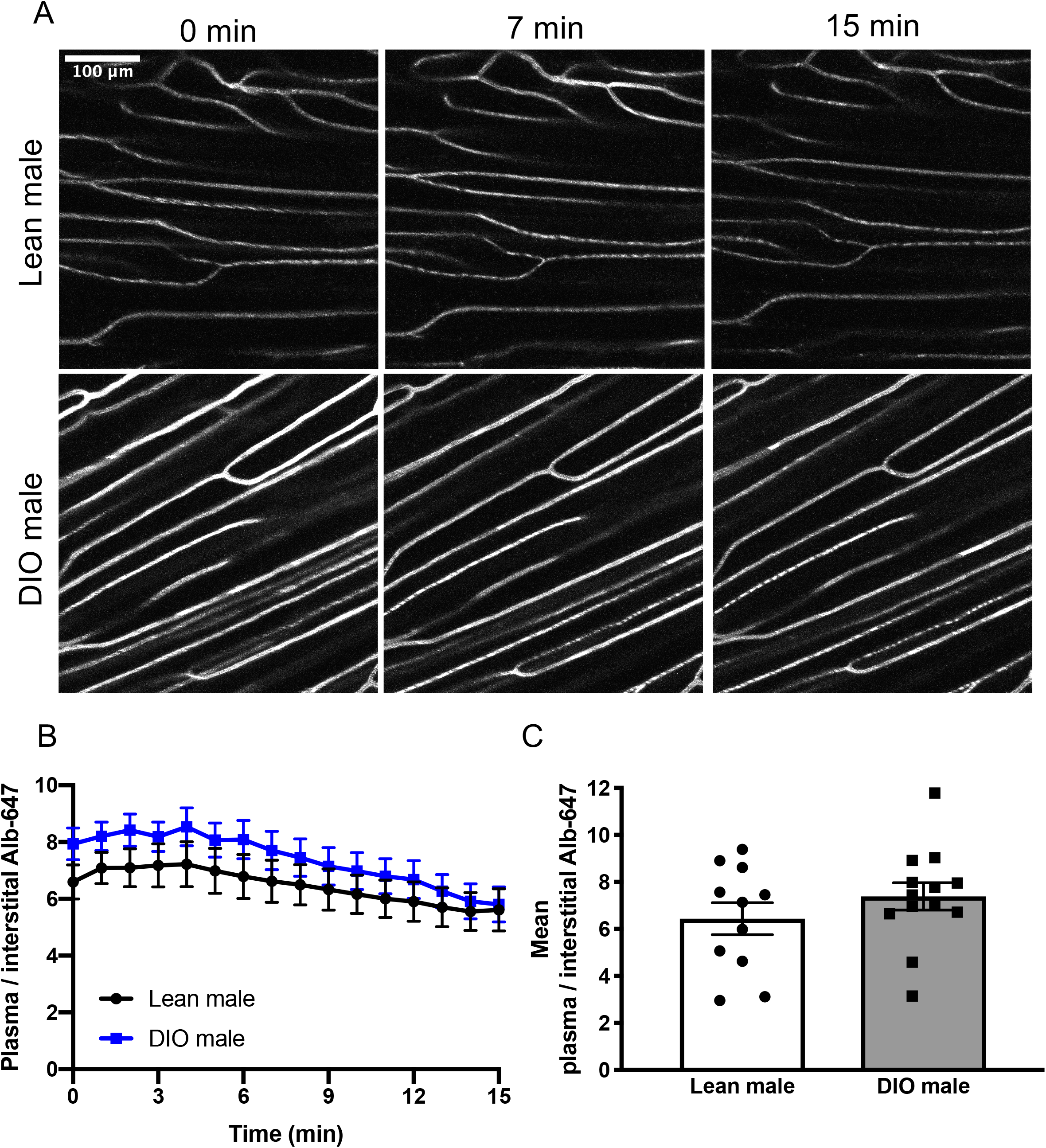
No difference in albumin equilibration between lean and DIO male mice. **A)** Representative Alb-647 images (maximum intensity projections) in lean (n=11) and DIO (n=13) male mice. **B)** The ratio of plasma to interstitial Alb-647 as a function of time following Alb-647 injection. **C)** Mean plasma to interstitial Alb-647 ratio over the course of the experiment. Alb-647 – albumin-647. Groups were compared using Student’s t-test.

### HFD does not alter SkM capillary endothelial ultrastructure nor trans-endothelial insulin transport in female mice

We next tested whether the effects of HFD on endothelial ultrastructure and EIT were similar in female mice. Female mice fed a HFD for 16 weeks weighed 30% more than their chow-fed counterparts (**Supplemental Figure 5A**). Interestingly, HFD-fed females did not have a significantly higher percentage of body fat than chow-fed mice (**Supplemental Figure 5B**).

We did not observe any difference in endothelial vesicular volume between chow and HFD-fed female mice (**Figure 7A&B**). Interestingly, the distribution of capillary vesicular densities appeared to shift from a normal distribution in chow-fed females to a more bimodal distribution in HFD-fed females (**Figure 7C**). This subtle shift in capillary vesicular distribution may represent an early stage of HFD-induced endothelial dysfunction. In general, however, female mice are largely protected from the deleterious effects of HFD on capillary endothelial ultrastructure.

**Figure 7:**
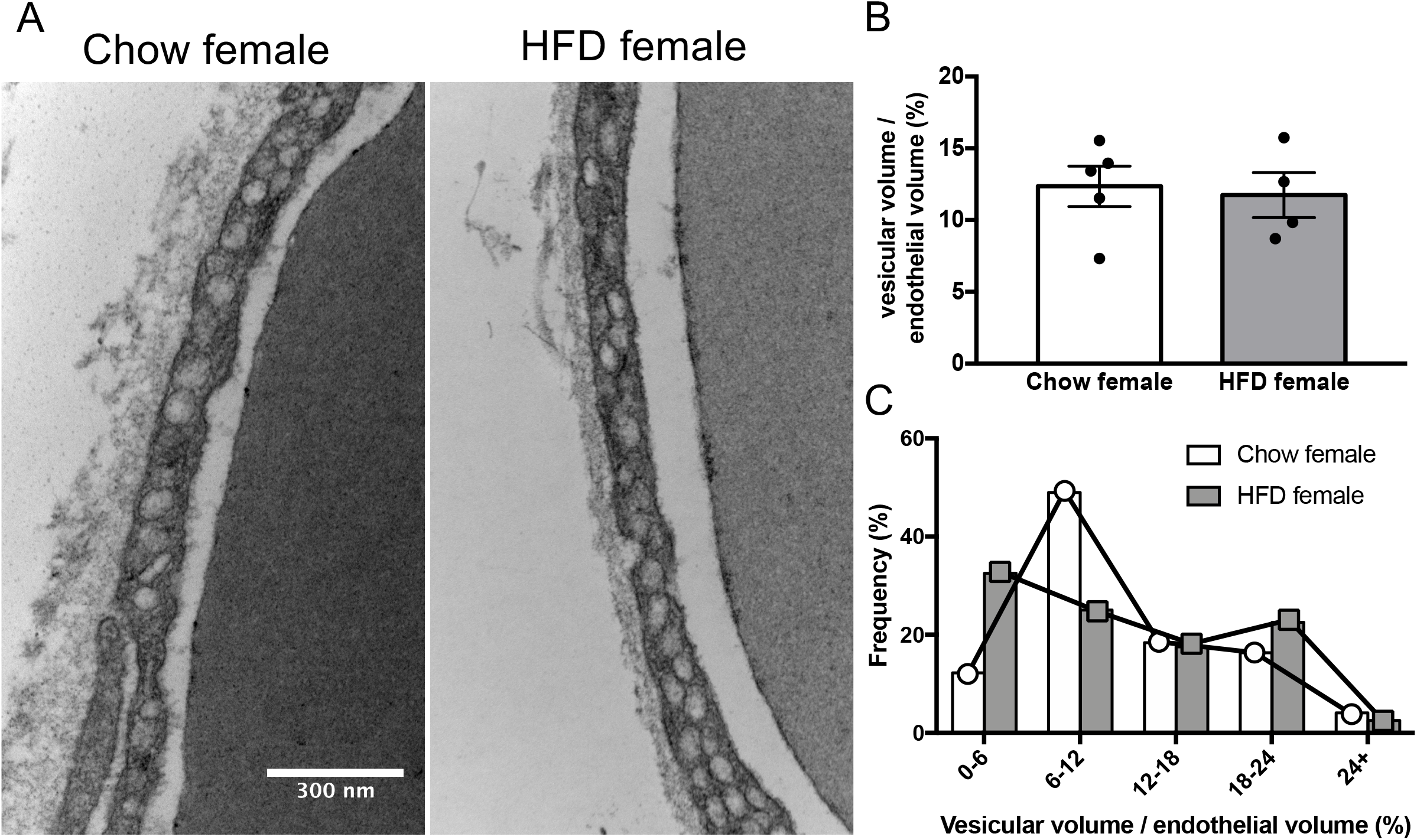
No effect of HFD on endothelial vesicles in female mice. **A)** Representative electron micrographs of the capillary endothelium in the gastrocnemius of chow and HFD-fed female mice. **B)** Volume of vesicles relative to total endothelial volume in chow (n=5) and HFD-fed (n=4) female mice. **C)** Frequency distribution of relative vesicular volume in all capillaries pooled from chow (n=49) and HFD-fed female mice (n=40). Groups were compared by Student’s t-test.

Given that endothelial ultrastructure is not affected by HFD in female mice, we hypothesized that EIT would also be unchanged in these mice. Indeed, we did not observe any difference in EIT between chow and HFD-fed females (**Figure 8**). Specifically, there was no change in the dissipation of the plasma/interstitial INS-647 gradient (**Figure 8B&C**). Similar to HFD-fed males, plasma INS-647 levels were higher in HFD-fed females following the INS-647 bolus (**Supplemental Figure 6A&B**). However, the rate of INS-647 clearance from the plasma was not different between the two groups (**Supplemental Figure 6A&B**). HFD-females tended to have higher levels of interstitial INS-647 (**Supplemental Figure 6C**) as expected from their higher levels of plasma INS-647. However, there was no difference in the rate of interstitial INS-647 appearance (**Supplemental Figure 6D**) between chow and HFD-fed females. With respect to microvascular hemodynamic indices, we did not observe any difference in either average plasma-perfused surface area (**Supplemental Figure 7A&B**) or capillary diameter (**Supplemental Figure 7C&D**). As expected from the lack of differences in capillary surface area (**Supplemental Figure 7B**) and EIT (**Figure 8C**), there was no change in total extravascular INS-647 delivery (**Supplemental Figure 6E&F**) in HFD-fed females. Following the intravital microscopy experiments, we did not observe any difference in terminal blood glucose (**Supplemental Figure 8A**). HFD-fed females did have a 27% non-significant reduction in 2[^14^C]DGP accumulation in the gastrocnemius (**Supplemental Figure 8B**). These findings suggest that, in contrast to males, 16 weeks of HFD neither impedes EIT nor causes SkM IR in female mice.

**Figure 8:**
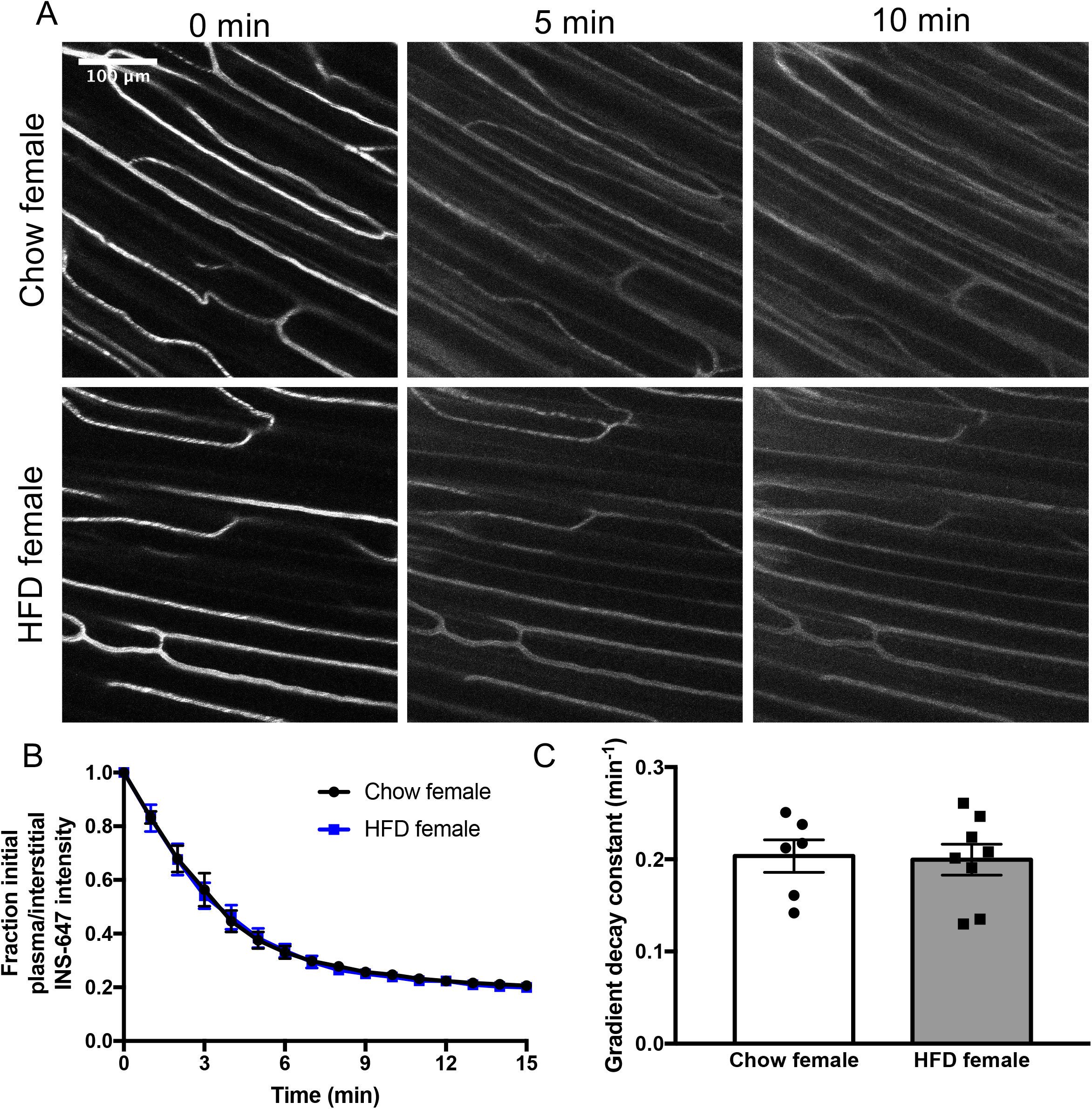
HFD does not alter trans-endothelial insulin transport in females. **A)** Representative INS-647 images (maximum intensity projections) in chow (n=6) and HFD-fed (n=8) female mice. **B)** The ratio of plasma to interstitial INS-647 as a function of time following INS-647 injection, normalized to the ratio at t = 0 min. **C)** The gradient decay constant of the plasma to interstitial INS-647 ratio as a function of time. Groups were compared using Student’s t-test.

## DISCUSSION

In this study, we found that 16 weeks of HFD causes a profound reduction in the number of endothelial vesicles in SkM capillaries in male mice. This ultrastructural damage to the endothelium is associated with impaired EIT *in vivo*, as determined by intravital microscopy. Of note, this is the first study to directly demonstrate impaired EIT in a mouse model of obesity and SkM IR. Interestingly, however, interstitial insulin delivery is maintained in DIO male mice through elevated plasma insulin. The fact that DIO male mice still show severe SkM IR even in the absence of impaired interstitial insulin delivery indicates that myocytes are the major contributor to SkM IR in this model. Female mice, on the other hand, were protected from the deleterious effects of HFD on endothelial ultrastructure, EIT, and SkM insulin sensitivity. In summary, these findings indicate that while HFD impairs EIT in male mice, interstitial insulin delivery can be sustained through hyperinsulinemia.

Most previous studies, but not all (21), have demonstrated that the delivery of insulin to SkM interstitial space is reduced in obese humans (18) and animal models (19, 20, 33). The delivery of insulin to SkM depends on both the capillary surface area for insulin exchange and the rate at which insulin moves across the capillary endothelium. Many studies have shown that insulin-stimulated SkM perfusion is reduced in obesity (2, 19, 34, 35). The contribution of EIT to the reduced insulin delivery observed in obesity, however, has been unclear due to an inability to make direct measurements *in vivo*. We recently developed an intravital microscopy technique to directly visualize and quantify EIT in individual SkM capillaries *in vivo* (6). This method has major advantages over *in vitro* studies of cultured ECs. Namely, cultured ECs are very different than capillaries *in vivo* with respect to microenvironment, protein expression, permeability, and the mechanism of insulin transport (6, 36–38). Utilizing direct measurements of EIT *in vivo*, we demonstrate that the movement of insulin across the endothelium is impaired in SkM capillaries of DIO male mice.

We observed a 15% reduction in EIT in DIO male mice. The implication of this reduction in EIT would presumably be a reduction in total delivery of insulin to SkM. Conversely, we found that interstitial insulin levels were actually elevated DIO male mice. One potential limitation of the high resolution intravital microscopy measurements is that they are made in a very small volume and may not reflect insulin delivery at the whole muscle level. Regardless, this finding emphasizes that, in addition to surface area for insulin exchange and EIT, plasma insulin concentration is an important determinant of muscle insulin delivery. In the current study, higher plasma insulin levels in DIO male mice compensate for the reduced EIT to maintain extravascular insulin delivery. Plasma insulin levels were most likely higher in DIO mice because the dose of INS-647 was normalized to total body mass and plasma volume does not scale with body mass in obese mice (39). That is to say, the plasma insulin concentration was likely higher in DIO male mice because more insulin was administered into a similar plasma volume. Nonetheless, the higher levels of plasma insulin alone would not be expected to affect EIT because 1) insulin efflux is not saturable (6) and 2) HFD-female mice also have higher levels of plasma insulin but no change in EIT. Furthermore, these experimental conditions recapitulate the hyperinsulinemia observed in obesity. Despite the higher levels of extravascular insulin in the DIO male mice, they still displayed SkM IR. This indicates cellular IR at the level of the myocyte. Thus, myocellular IR can be compounded by reduced EIT to severely impair insulin action in SkM during obesity.

The reduction of EIT in DIO male mice is associated with a profound reduction in the density of endothelial vesicles. This is consistent with a previous finding from Bagi and colleagues who showed that the number of endothelial vesicles in coronary arterioles was reduced in humans with established Type 2 diabetes (40). Endothelial vesicles can serve as macromolecular transporters (31) which may carry insulin (41). It is unclear whether these vesicles are distinct, transcytotic carriers or if they represent pores in the endothelium (42). We hypothesize that these capillary endothelial vesicles are likely components of pore networks for two reasons. First, we previously showed that trans-endothelial insulin transport is not receptor-mediated nor saturable *in vivo* (6). We have further validated these findings in this study by showing that the majority of SkM ECs do not express the insulin receptor. Assuming that endothelial vesicles do indeed carry insulin (41), the transport of insulin through endothelial pores is more compatible with the non-saturable nature of insulin transport. The presence of a non-saturable yet active, energy-requiring transcytotic transport mechanism seems unlikely. Furthermore, if individual vesicles were actively taking up macromolecules and being transcytosed across the endothelium, we would expect to see molecular accumulation in the capillary endothelium. Unlike studies utilizing cultured ECs (43), we have never observed the accumulation of either INS-647 or rho-dex in the lining of SkM capillaries. It is possible that vesicular transcytosis occurs so rapidly in capillaries as to preclude endothelial accumulation. We did, however, observe accumulation of these molecules in the cells lining venules, indicating that vesicular transcytosis may be a more relevant transport mechanism in larger blood vessels. The role of venular insulin transport in SkM insulin action is currently unknown and warrants further investigation. In summary, we hypothesize that, in DIO male mice, the number of capillary endothelial pores available for insulin exchange is reduced, thereby impairing EIT.

It should be noted that the magnitude in reduction of endothelial vesicles (45%) was much larger than the impairment in EIT (15%). We have three hypotheses for this discrepancy. First, it is possible that paracellular permeability, as governed by the width of inter-endothelial junctions, increases in DIO male mice to compensate for the reduction in transcellular (i.e. vesicular pores) permeability. In this case, one would expect that a significant increase in paracellular permeability would be reflected by an increase in capillary albumin. However, transendothelial albumin equilibration was the same in lean and DIO male mice. Thus, it is unlikely that there is significant compensatory paracellular transport in DIO male mice. A more likely explanation is that there is an excess of endothelial pore surface area available for insulin exchange in healthy, lean mice. Thus, a more significant reduction in endothelial vesicular density may be required to severely limit EIT. Finally, we cannot exclude the possibility that the paracellular movement of insulin through inter-endothelial junctions is a greater determinant of EIT than trans-cellular insulin movement.

Our intravital microscopy technique also allows for the measurement of indices of microvascular structure and hemodynamics. We did not observe any difference in plasma-perfused surface area and capillary diameter between lean and DIO male. Furthermore, the administration of insulin did not affect plasma-perfused surface area. This suggests that there was no effect of HFD on the capillary surface area available for insulin exchange. These findings are seemingly in contrast to previous studies utilizing contrast-enhanced ultrasound which concluded that 1) insulin increases SkM perfusion and 2) insulin-stimulated SkM perfusion is reduced in obesity (5, 19). The use of contrast-enhanced ultrasound to measure the distribution volume of ~1-3μm microbubbles gives an index of red blood cell (RBC) distribution volume. This is different from our measurement of the distribution volume of a rhodamine-labeled plasma marker, 2-megadalton dextran. As insulin exists in plasma and not RBCs, the measurement of plasma-perfused surface area is more relevant for assessing the surface area available for insulin exchange. Thus, it is possible that insulin and HFD have divergent effects on RBC and plasma distribution. Namely, while insulin may increase and obesity may decrease microbubble distribution volume (5), neither insulin nor obesity alter plasma-perfused surface area in SkM.

In contrast to male mice, we found that female mice were protected from the deleterious effects of HFD on EIT and endothelial ultrastructure. The most likely explanation for this finding is that, while HFD-fed female mice were heavier than chow-fed mice, they did not have a significantly higher percentage of body fat. This observation is mostly consistent with a previous study which showed that 15 weeks of HFD causes weight gain but only a small increase in body fat percentage in female mice (44). Thus, HFD in the absence of obesity does not cause impaired capillary insulin transport in female mice. Consistent with the unaltered EIT, SkM insulin sensitivity, as measured by 2[^14^C]DGP accumulation, was only slightly and non-significantly reduced in HFD-fed females. Moreover, endothelial vesicular density was not altered by HFD in females. This contrasts with male mice which developed severe SkM IR and showed a dramatic reduction in endothelial vesicular density when fed a HFD. It is also conceivable that sex differences in microvascular function (45, 46) protect female mice from the endothelial dysfunction associated with HFD.

In summary, we have shown for the first time that the movement of insulin across the endothelium in SkM capillaries is impaired in a mouse model of diet-induced obesity and IR. This impairment in EIT is associated with profound alterations to endothelial ultrastructure. These findings demonstrate that capillaries limit the delivery of insulin to insulin resistant skeletal myocytes. Potential strategies to accelerate EIT include maintaining endothelial ultrastructure, widening inter-endothelial junctions, or increasing capillary surface area for exchange. These approaches may be useful in enhancing the delivery of insulin to SkM for the treatment of IR or in generating fast-acting insulin analogs for diabetics.

## Acknowledgments

The authors thank Freyja James (Department of Molecular Physiology and Biophysics, Vanderbilt University) for performing catheterization surgeries. The authors gratefully acknowledge the use of services provided by the Vanderbilt Cell Imaging Shared Resource, Mouse Metabolic Phenotyping Center, Research Electron Microscopy Facility, and the Diabetes Research and Training Center.

## Author contributions

The project was conceived and designed by IMW and DHW. IMW, JSB, DPB, and FAV performed experiments and collected data. IMW wrote the manuscript. Manuscript edits and discussions were contributed by PMM, JSB, FAV, and DHW. IMW is the guarantor of this work and, as such, had access to all data and takes full responsibility for the work as a whole.

## Prior Presentation

Parts of this study were presented at the American Diabetes Association Meeting in San Diego, CA, June 2017.

