## Supplemental Figures for "Trans-Endothelial Insulin Transport is Impaired in Skeletal Muscle Capillaries of Obese Male Mice"

### Figure 1

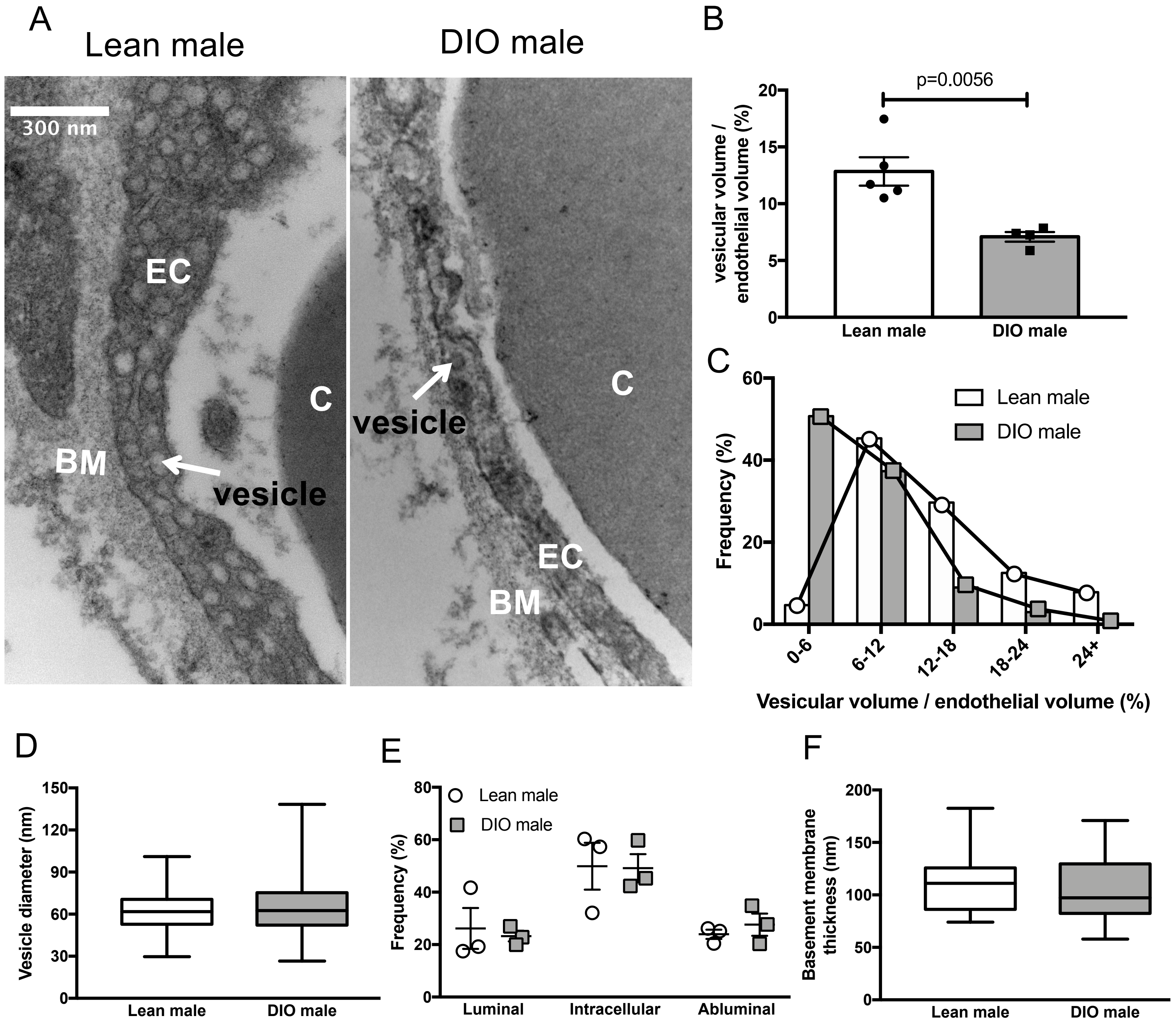


**Figure 1: Skeletal muscle capillaries of DIO male mice contain fewer endothelial vesicles. A)** Representative electron micrographs of the capillary endothelium in the gastrocnemius of lean and DIO male mice. **B)** Volume of vesicles relative to total endothelial volume in lean (n=5) and DIO (n=4) male mice. **C)** Frequency distribution of relative vesicular volume in all capillaries grouped from lean (n=64) and DIO male mice (n=66). **D)** The average diameter of all endothelial vesicles in lean (n=227) and DIO (n=116) male mice. **E)** Frequency distribution of the localization of vesicles in the capillary endothelium. **F)** Average basement membrane thickness in capillaries from lean (n=31) and DIO male mice (n=31). In the box and whisker blots, the box extends from the 25$^{th}$ to the 75$^{th}$ percentiles and the whiskers indicate the range. Groups were compared using Student's t-test. C – capillary lumen, EC – endothelial cell, BM – basement membrane.

### Figure 2

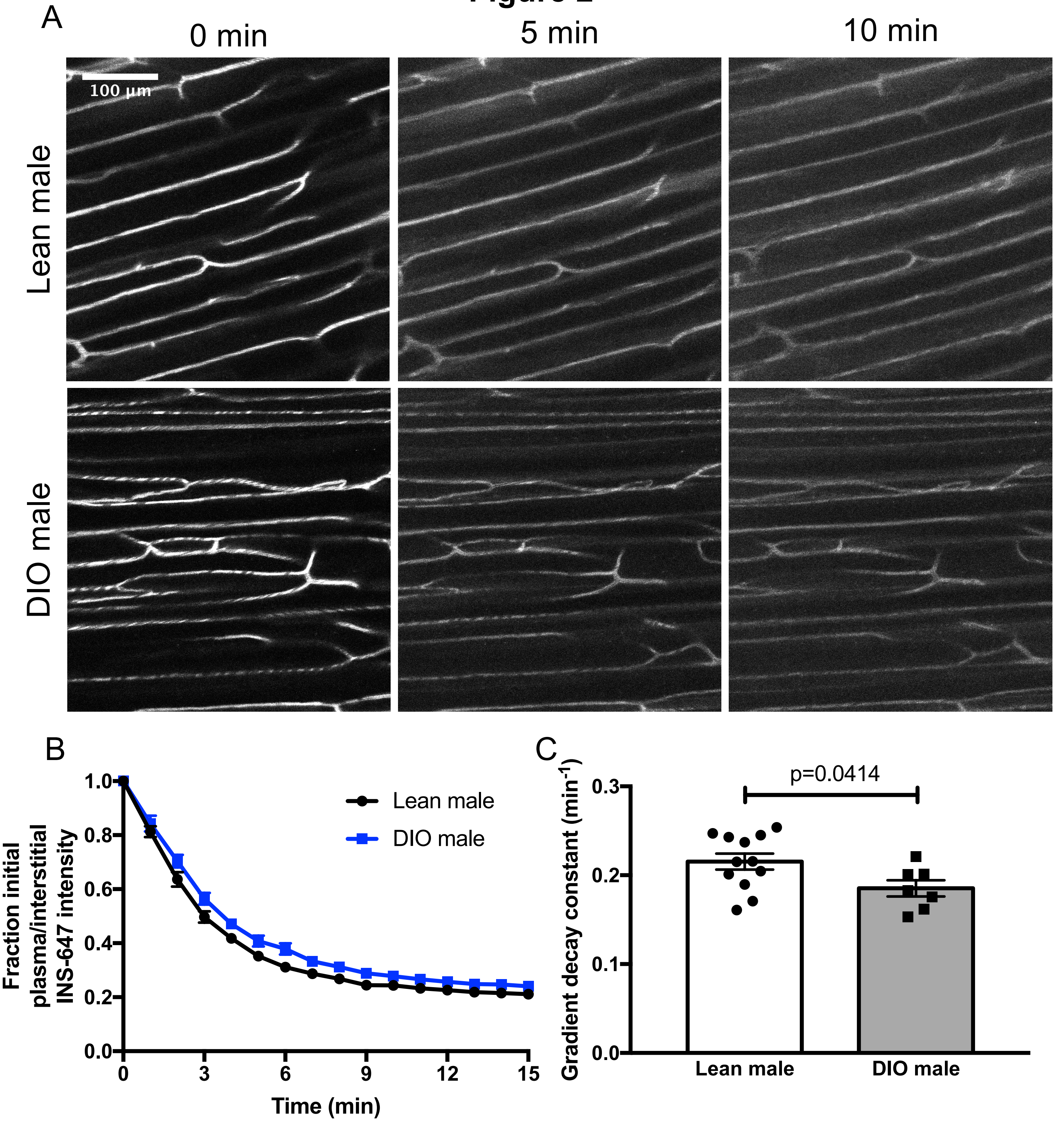


**Figure 2: Obese male mice have impaired trans-endothelial insulin transport in skeletal muscle capillaries. A)** Representative INS-647 images (maximum intensity projections) in lean (n=12) and DIO (n=7) male mice. **B)** The ratio of plasma to interstitial INS-647 as a function of time following INS-647 injection, normalized to the ratio at t = 0 min. **C)** Decay constant of the plasma / interstitial INS-647 ratio, a measure of trans-endothelial insulin transport kinetics. INS-647 – insulin-647, DIO – diet-induced obese. Groups were compared using Student's t-test.

### Figure 3

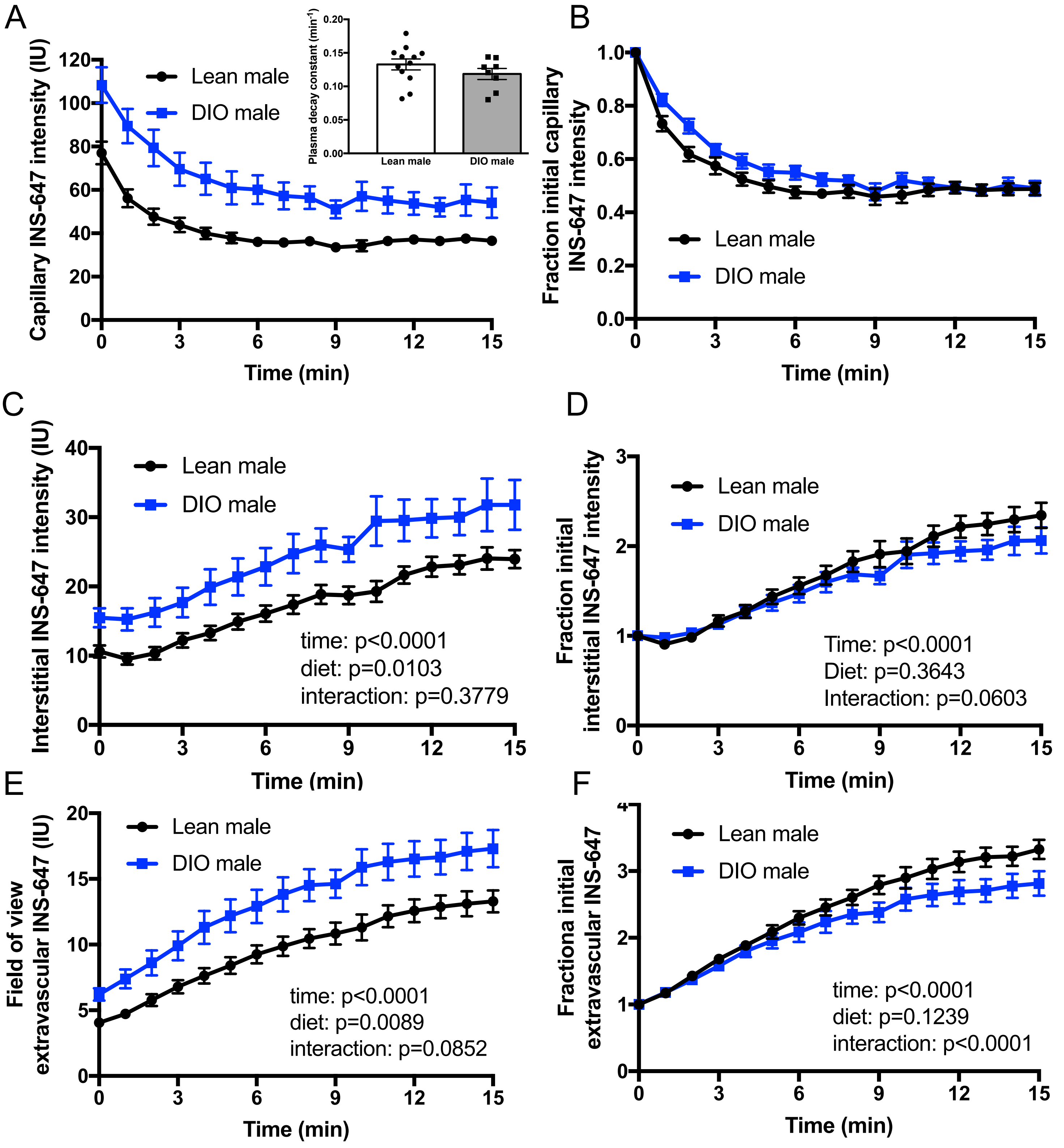


**Figure 3: Effects of HFD on plasma insulin clearance, interstitial appearance, and insulin delivery in male mice. A)** Capillary plasma INS-647 intensity as a function of time following injection in lean (n=12) and DIO (n=7) male mice. Inset shows the decay constant of capillary INS-647. **B)** Data in **A** normalized to the capillary INS-647 intensity at t = 0 min. **C)** Interstitial INS-647 intensity as a function of time following injection. The interstitial space is defined as the region emanating 1-3μm from the capillary wall. **D)** Data in **C** normalized to the interstitial INS-647 intensity at t = 0 min. **E)** Total extravascular INS-647 in the field of view as a function of time following injection. **F)** Data in **E** normalized to the extravascular INS-647 intensity at t = 0 min. Groups were compared either by Student's t-test or by two-way repeated measures ANOVA. IU – intensity units.

### Figure 4

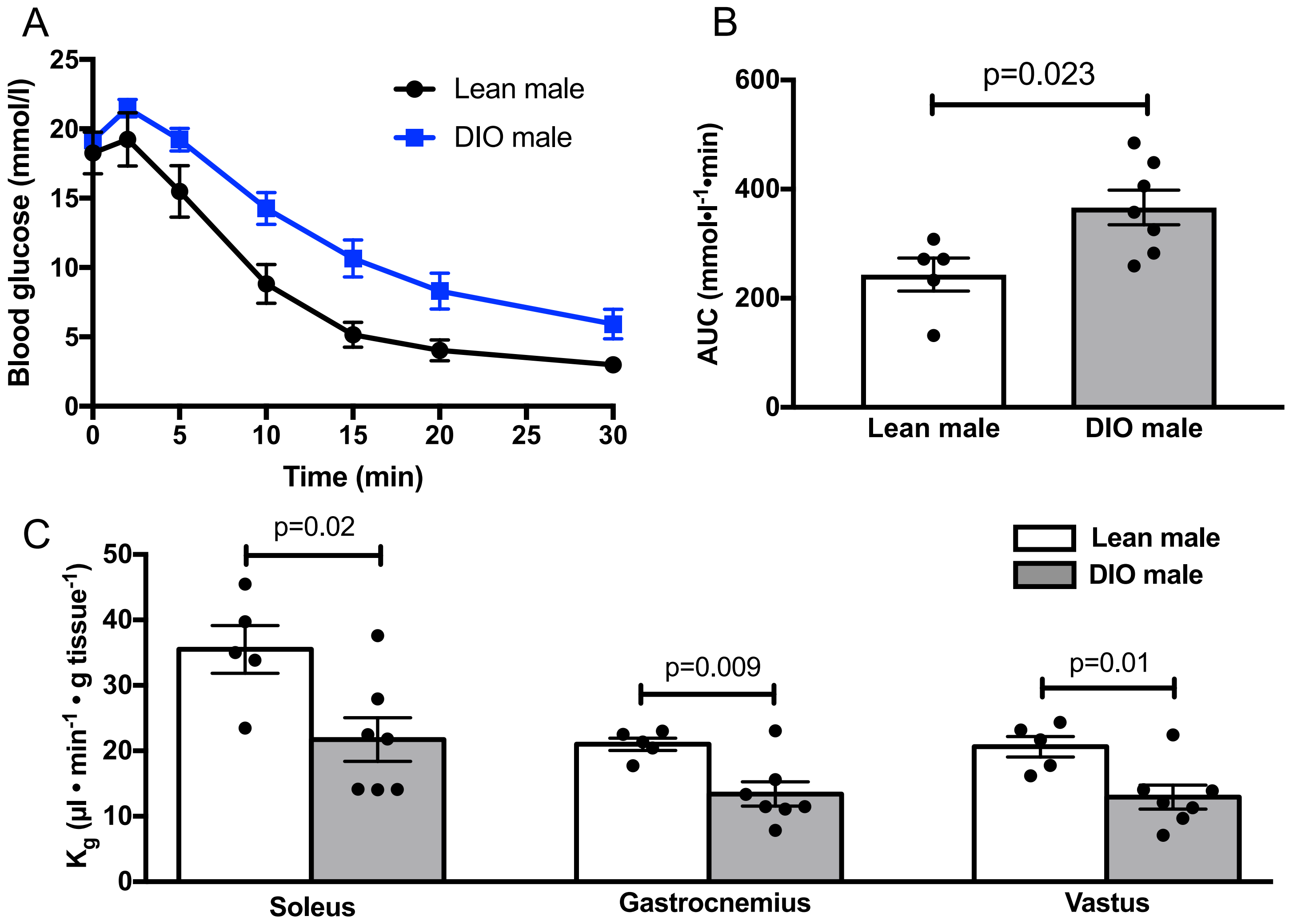


**Figure 4: DIO male mice display skeletal muscle insulin resistance. A)** Glucose excursions in anesthetized lean (n=5) and DIO (n=7) male mice following a 4U/kg intravenous insulin bolus. **B)** Area under the glucose excursion curves in **A**. **C)** Clearance of 2[$^{14}$C]2deoxyglucose during the insulin tolerance tests by the soleus, gastrocnemius, and vastus muscles. Groups were compared using Student's t-test.

### Figure 5

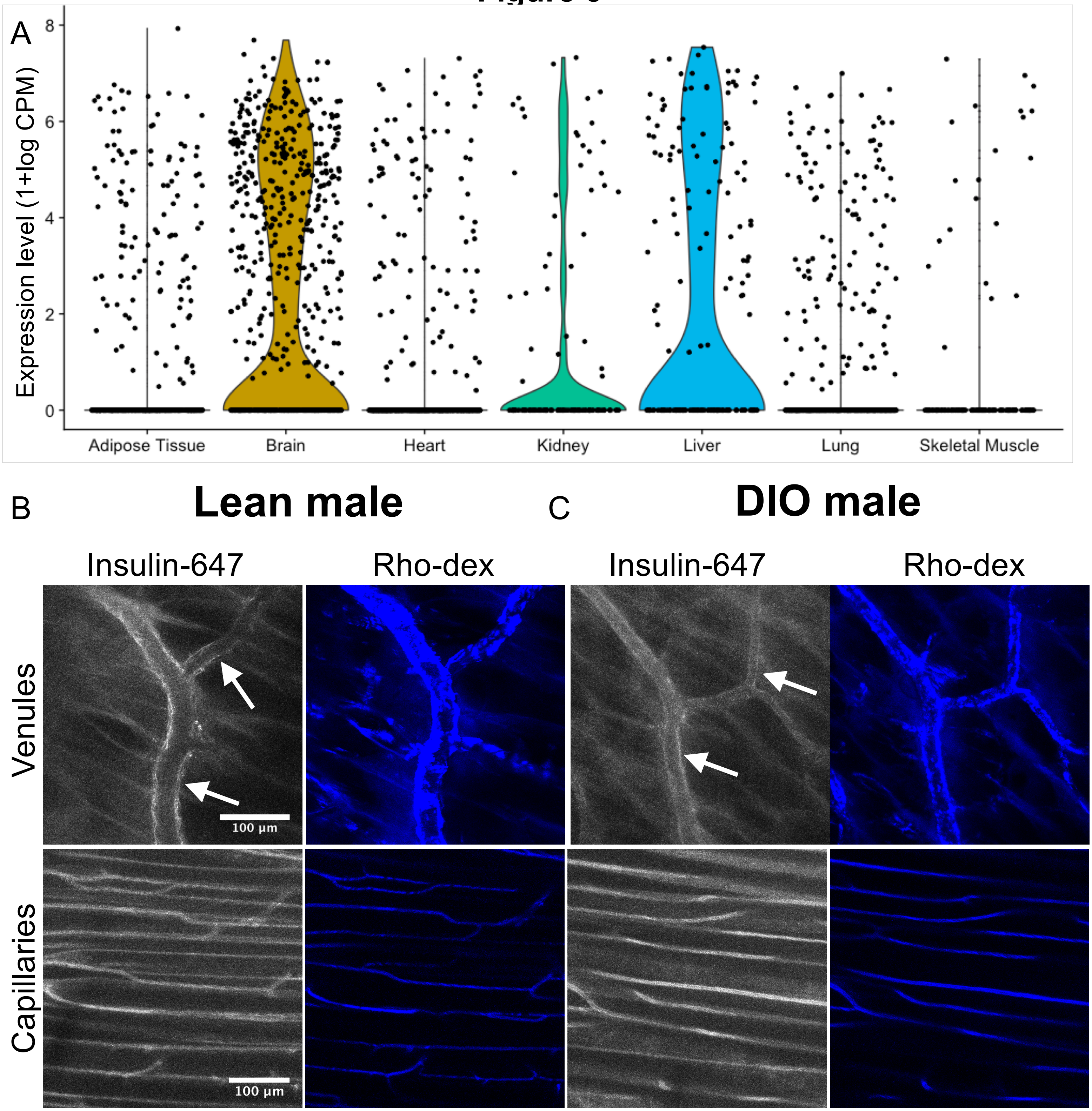


**Figure 5: Capillary trans-endothelial insulin transport in skeletal muscle does not involve the insulin receptor or endothelial insulin accumulation. A)** Violin plots of insulin receptor mRNA expression in tissue-specific endothelial cells as determined by single-cell RNA sequencing. **B,C)** Intravital microscopy images of insulin-647 and rhodamine-labeled 2MDa dextran (Rho-dex) in venules (top panels) and capillaries (bottom panels) from **B)** lean and **C)** DIO male mice. Both insulin and dextran can be seen accumulating in the endothelium of venules but not capillaries. Arrows indicate regions of insulin accumulation in the venular endothelium. Rho-dex – 2MDa tetramethylrhodamine-dextran, CPM – counts per million.

#### Figure 6

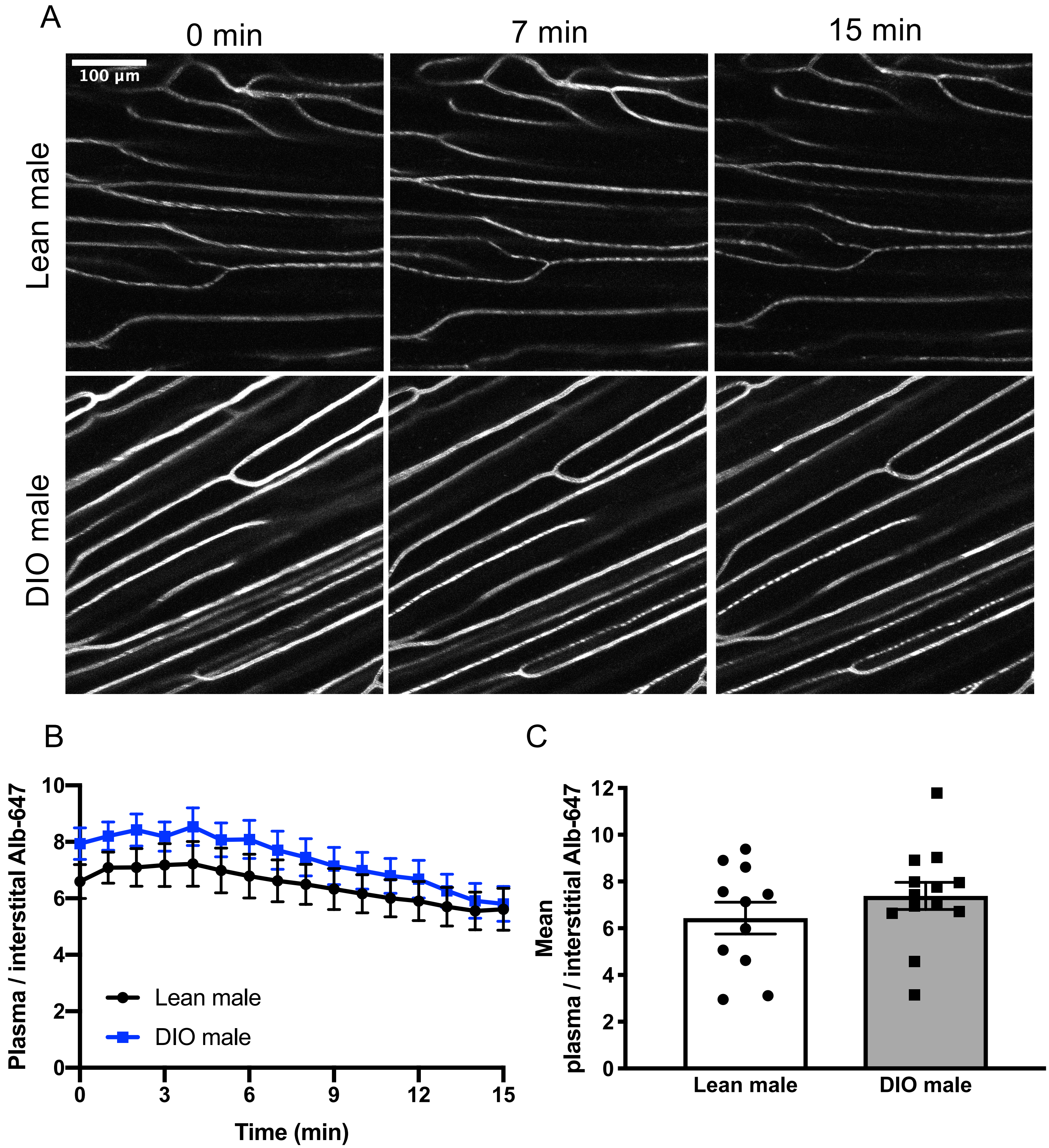


**Figure 6: No difference in albumin equilibration between lean and DIO male mice. A)** Representative Alb-647 images (maximum intensity projections) in lean (n=11) and DIO (n=13) male mice. **B)** The ratio of plasma to interstitial Alb-647 as a function of time following Alb-647 injection. **C)** Mean plasma to interstitial Alb-647 ratio over the course of the experiment. Alb-647 – albumin-647. Groups were compared using Student's t-test.

### Figure 7

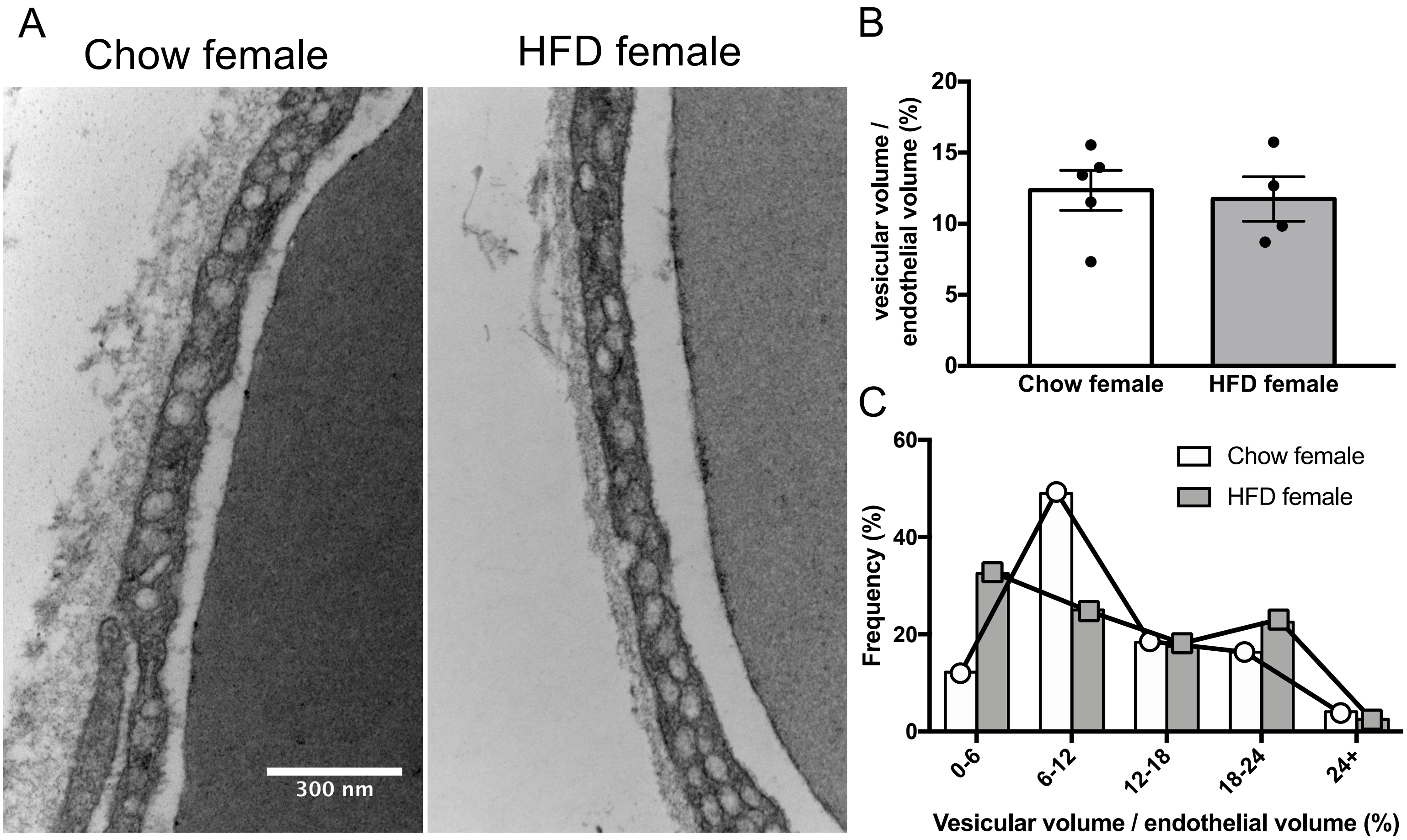


**Figure 7: No effect of HFD on endothelial vesicles in female mice. A)** Representative electron micrographs of the capillary endothelium in the gastrocnemius of chow and HFD-fed female mice. **B)** Volume of vesicles relative to total endothelial volume in chow (n=5) and HFD-fed (n=4) female mice. **C)** Frequency distribution of relative vesicular volume in all capillaries pooled from chow (n=49) and HFD-fed female mice (n=40). Groups were compared by Student's t-test.

### Figure 8

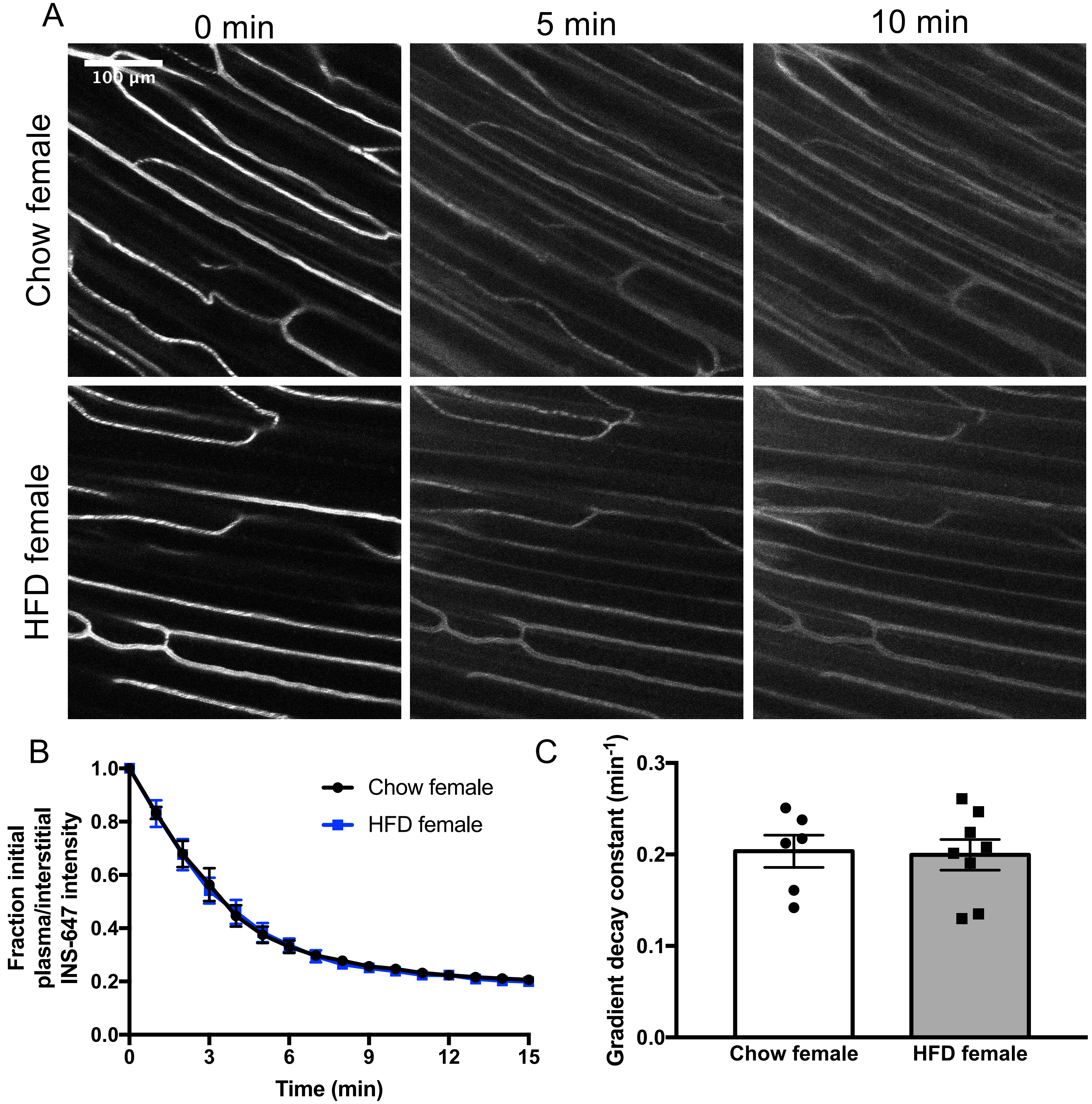


**Figure 8: HFD does not alter trans-endothelial insulin transport in females. A)** Representative INS-647 images (maximum intensity projections) in chow (n=6) and HFD-fed (n=8) female mice. **B)** The ratio of plasma to interstitial INS-647 as a function of time following INS-647 injection, normalized to the ratio at t = 0 min. **C)** The gradient decay constant of the plasma to interstitial INS-647 ratio as a function of time. Groups were compared using Student's t-test.
